# Sorry, we’re open: Golden Open Access and inequality in the natural sciences

**DOI:** 10.1101/2020.03.12.988493

**Authors:** Russell J Gray

## Abstract

Global Open Access (GOA) journals make research more accessible and therefore more citable; however, the publication fees associated with GOA journals can be costly and therefore not a viable option for many researchers seeking high-impact publication outlets. In this study, I collect metadata from 237 open-access natural science journals and analyze them in terms of Article Processing Charges (APC), Impact Factor (IF), Eigen Factor (EF), citability, and country of publisher. The results of this study provide evidence that with IF, EF, and citability all increase as APC increases, and each of these metrics are higher in publishers from developed countries in comparison to developing countries. Implications of these trends are discussed in regards to natural sciences and inequality within the global scientific community.

## Introduction

With the prominence of entities such as Sci-Hub (Black Open Access) and ResearchGate (Green Open Access), the availability of journal articles has increased and all but nullified the previous paywall system from creating capital for journals (Fuchs and Sandoval, 2013). While this increase in accessibility of scientific literature has removed constraints of researchers and the general public on a global scale, it has also caused many high-impact journals to transition to mandatory open access, also known as Golden Open Access (GOA), in order to continue their profitability through scientific publication (Piwowar, et al. 2018; Lewis, 2012; Harnad, et al. 2008). Although the concept of open-access science may seem genuine on its face, the cost associated with publishing in open-access journals, and through this cost, perceived relaibility and importance, is what makes this system problematic.

Although many journals enable funding assistance for certain countries and institutions, and Article Processing Charges (APC) can sometimes be factor into project funding, this isn’t the case for all researchers and projects, especially when it comes to more expensive, prestigious journals. A study by Ellers et al. (2017) provided evidence that researchers from institutions in developing countries are still paying the full price of developed western nations to publish in high-impact “Mega-Journals” in order to gain exposure and credibility for their research. Unfortunately, the alternative, especially when it comes to open-access, is to publish in journals with a lower Impact Factor (IF); otherwise, the author publish in predatory journals which have low-quality peer review, or publish other questionable articles which then impact the credibility of a manuscript by affiliation (Beall, 2013).

In the current system of academic publishing, the publication process can be an incredibly stressful for researchers, and in many cases time-consuming (Björk and Solomon 2013). One of the most important benefits of high-impact journals are their ability to generate exposure through increased citations. Since high-impact journals have built a reputation of credibility, the review process, and therefore the content of the publication is not questioned as much as a non-reputable journal with a lower impact-factor, which can cause immediate and noticeable acceleration in the careers of researchers (Reich, 2013). However, the prices of high-impact GOA journals become problematic for researchers without available funding to publish, because options to make potentially important research impactful become more and more narrow.

In this study, I analyze metadata of open-access journals to determine links between publication costs, impact factor, eigen factor, and citability for journals pertaining to the natural sciences. I then discuss the implications of inequality among the global scientific community in regards to the open-access framework and its costly limitations.

## Methods

Data from the Directory of Open Access Journals (Morrison, 2008) was used to acquire journal name, APC, country of publisher, and currency records (n = 14275). R statistical software was used for all data analyses; packages used in the analyses included: 1) the scholar package (Keirstead, 2016) to generate impact factors using journal name strings; 2) the quantmod package (Ryan et al. 2019) to convert all currencies to USD for standardization; 3) the dplyr package (Wickham et al. 2015) and reshape2 package (Wickham, 2012) to clean and select specific data, and 4) ggplot2 package (Wickham et al. 2016) for data visualizations.

A subset of journals (n = 1047) which had topics relative to the biological sciences was queried by creating a subset of keyword data from the original Directory of Open Access dataset (e.g. ecology, conservation, agriculture, species, forest, etc.). Large disparities between natural sciences and medical science in relation to impact factor, so to further clean the data, journals with an IF >40 were excluded from the analysis. The excluded journals were checked to via keywords to ensure none were related to the natural sciences; excluded journals were related to oncology, medical, and engineering. The final analyses were performed on a deeply vetted dataset of open-access, natural science journals (n = 237).

In order to explore potential trends in the data, APC was examined in relation to citability (total number of citations), IF (general impact of the journal), and Eigen Factor (relative importance of the journal). The country of each journal publisher was also examined against cost, citability, IF, and Eigen Factor (EF hereafter) to locate any potential disparities between them. Using functions from the ggpubr package (Kassambara, 2020), Pearson’s correlation coefficient was used to analyze potential correlation between APC, IF, EF, and citability of journals. Country of publisher was also examined in order to determine if there were any trends between country and APC, IF, EG, and citability.

## Results

A linear trend was found between IF, EF, and citability in relation to APC. Additionally, it was found that journal publishers from developed countries were more likely to have higher APC, IF, EF, and citability (Figure 1). Mean APC of open-access journals analyzed (n = 237) was $1344.27 USD, mean IF was 2.855, mean EF was 0.017, and mean total citation count or citability was 5925.92. APC was found to be significantly positively correlated with IF (Pearson correlation: r(237) = 0.391, p < .001). There was also a highly significant positive correlation between APC and EF (Pearson correlation: r(237) = 0.303, p < .001). Finally, there was a highly significant positive correlation between APC and citability (Pearson correlation: r(237) = 0.323, p < .001).

**Figure 1.**
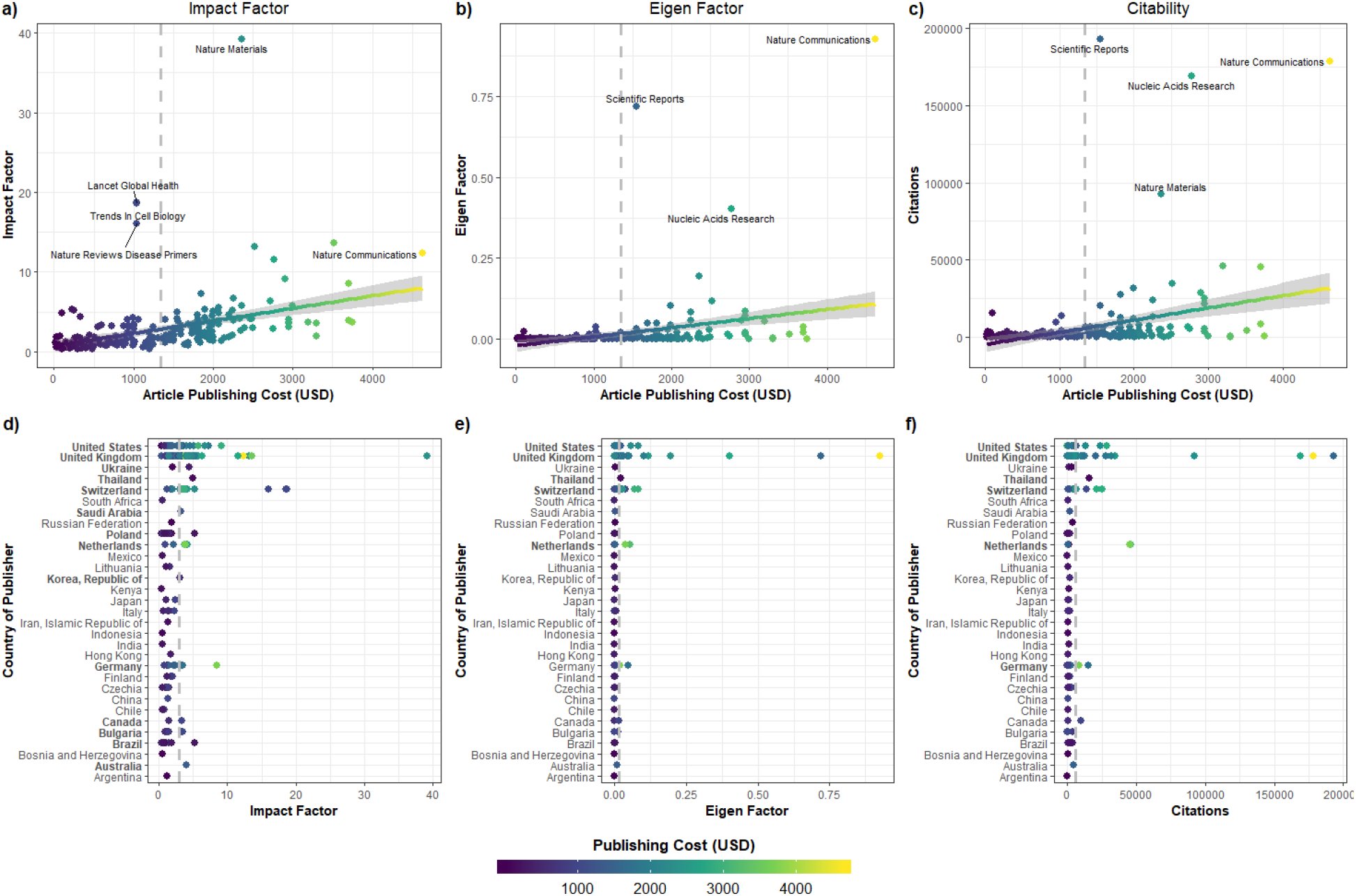
Trends in IF, EF, and citability: a) APC and IF, b) APC and EF, c) APC and total number of citations; for each of the three upper graphs, mean APC is represented as a dashed vertical line for each of the three upper graphs, a General Linear Model (GLM) trendline is fitted to the data with standard error shadow, and outliers are labeled with the journal name; d) IF of journals by country of publishers; e) EF of journals by country of publishers; f) citability of journals by country of publishers; for the three lower graphs, all publishing countries which exceed the mean IF, EF, and citation total (dashed vertical line) are represented in bold.

Journals published in the United States, United Kingdom, Ukrain, Thailand, Switzerland, Saudi Arabia, Polan, Netherlands, South Korea, Germany, Canada, Bulgaria, Brazil, and Australia were more likely to have an IF above the overall mean than undeveloped countries. Journals published in the United States, United Kingdom, Thailand, Switzerland, and the Netherlands were more likely to have an EF above the overall mean. Journals publish in the United States, United Kingdom, Thailand, Switzerland, Netherlands, and Germany were more likely to have more total citations than the mean citability of other nations. Publishing cost was highest journals from the United States, United Kingdom, Switzerland, Netherlands, and Germany.

## Discussion

The data in this study shows that with increasing publishing costs, the impact (IF), importance (EF), and overall number of citations in open-access natural science journals also increase. The data also shows that the majority of journal publishers with the highest APC, IF, EF, and citability are from developed countries, which indicates that research published in journals from developing countries is less likely to gain exposure, while journals from developing countries are less likely to generate profit and, more importantly, credibility.

Expensive APC in prestigious journals are not linked to reliability of the research published in them (Brembs, 2018). Additionally, the impact factor of prestigious journals also creates a false perception of reliability in research they have published (Brembs, 2013). The paradox of GOA journals is that the system itself is inherently flawed, where the more manuscripts a journal publishes, the more capital it stands to generate, which gives rise to predatory journals and editorial complacency (Beall 2013). Therefore, GOA journals may create an environment where publisher profit is prioritized over scientific integrity of published research, while still maintaining prestige as an outlet of highly-credible research over other journals.

Unfortunately, the irregular distribution of resources throughout the world, and correspondingly in academia, creates a bubble where higher ranking institutions have more access to expensive, higher impact journals, while lower ranking institutions are forced to publish in less expensive or closed access journals (Siler, et al. 2018). The alternative is that authors from developing countries have to pay the prices of developed countries in order to have access to higher impact journals (Ellers, 2017). The options become even more narrow for individual researchers and groups who have no academic institutional affiliation, whether it be by personal career choice, transitionary period, or numerous other situations that prevent the benefits of institutional finance allowances (Burchardt, 2014). This is increasingly more problematic for scientific output, as independent researchers with no institutional affiliation have been on the rise over the past decade (ElSabry, 2017). However, evidence suggests that the researchers being put on the backburner, are no less reliable than the privileged few institutions which have the advantages to publish in high impact, high APC journals (Brembs, 2018; Brembs, 2013; ElSabry, 2017; Siler, et al. 2018).

Although there are many financial support options for GOA journals, there are also many caveats to their eligibility criteria. For example, once an international collaborator from a developed country is named as an author on a manuscript of authors which would otherwise be eligible for publication funding assistance such as Research4Life (Research4Life, 2015), the eligibility for financial assistance becomes void. I know this from personal experience after being rejected from funding assistance multiple times while attempting to publishing important studies on the critically endangered Sumatran elephant in GOA journals. While my co-authors were from eligible developing countries, I am from the United States, but have no current institutional affiliation and therefore none of the accompanying financial benefits, and none of us had access to the >1000 USD for publication fees. Does this mean the research we were reporting is not important, or reliable? Not at all. However, our options were narrowed significantly down to low-impact journals, which consequently are cited much less, and generally seen as less credible sources of scientific information. Cases like this for others may force an author to be removed from a manuscript in order to gain access to publication funds, which is not an environment that the process of publishing scientific research should ever be responsible of creating; nor should potentially important research on conservation of critically endangered species fly under the academic radar due to APC funding constraints preventing research exposure.

Since the journals analyzed in this study are related to natural sciences, this results also have implications for the research regarding ecology and conservation of species. Most biodiversity hotspots are located in the tropics; therefore, the vast majority of conservation research is conducted in tropical regions (Myers et al 2000). While many nations in tropical regions fall within the criteria for APC assistance from Research4Life as they are considered developing or under-developed countries (Research4Life, 2020), the majority of research coming from these undeveloped tropical regions historically include authors from developed countries (Stocks, 2008). With the current criteria systems in place for APC funding assistance in GOA journals for natural sciences through Research4life, foreign authors from developed countries would nullify the eligibility for assistance with publication costs, forcing authors to seek closed-access or hybrid journals, low-impact open-access journals, or potentially predatory journals to publish their research.

To conclude, the current transition to GOA publishing by prestigious, high impact journals, namely in the natural sciences, shows trends in inequality in global research output. Inequality amongst institutions, publishers, journals, and researchers is likely to prevent adequate exposure of potentially important research, while promoting the false ideology that prestige and costliness of journal publications are the equivalent of reliable science (Brembs, 2018; Brembs, 2013; ElSabry, 2017; Siler, et al. 2018). In our current age of technology, most journals have flipped the publication model from print to digital and online (Beall, 2013; Fitzpatrick, 2011), and even with a decrease in hardcopy production, publication fees are still increasing (Morrison, 2018). Journal publications, research impact, and citations are the academic currency of career scientists and scholars (Hirsch, 2005) and the scientific community is international. We must address these issues to create a more inclusive environment for important research to be recognized and researchers to prosper on a global scale, regardless of country or institutional affiliation.

